# Enhanced neuronal regeneration in the CAST/Ei mouse strain is linked to expression of differentiation markers after injury

**DOI:** 10.1101/160366

**Authors:** Véronique Lisi, Bhagat Singh, Michel Giroux, Elmer Guzman, Michio W Painter, Yung-Chih Cheng, Eric Huebner, Giovanni Coppola, Michael Costigan, Clifford J. Woolf, Kenneth S Kosik

**Affiliations:** Neuroscience Research Institute, Department of Molecular, Cellular, and Developmental Biology, University of California, Santa Barbara, Santa Barbara, CA 93106, USA; F.M. Kirby Neurobiology Center, Boston Children’s Hospital and Harvard Medical School, Boston, MA 02115, USA; Departments of Psychiatry and Neurology, Semel Institute for Neuroscience and Human Behavior, David Geffen School of Medicine, University of California, Los Angeles, Los Angeles, CA 90095, USA; Anaesthesia Department, Boston Children’s Hospital and Harvard Medical School, Boston, MA 02115, USA

## Abstract

Peripheral nerve regeneration after injury requires a broad program of transcriptional changes. We investigated the basis for the enhanced nerve regenerative capacity of the CAST/Ei mouse strain relative to C57BL/6 mice. RNA sequencing of dorsal root ganglia (DRG) showed a CAST/Ei specific upregulation of *Ascl1* after injury. *Ascl1* overexpression in C57BL/6 mice DRG neurons enhanced their neurite outgrowth. *Ascl1* is regulated by miR-7048-3p, which is down-regulated in CAST/Ei mice. Inhibition of miR-7048-3p enhances neurite outgrowth. Following injury, CAST/Ei neurons largely retained their mature neuronal profile as determined by single cell RNAseq, whereas the C57BL/6 neurons acquired an immature profile. These findings suggest that one facet of the enhanced regenerative phenotype is preservation of neuronal identity in response to injury.

## Introduction

Axonal regeneration after injury involves cell intrinsic and extrinsic factors. Insufficient information is currently available about the molecular mechanisms driving regeneration to design therapeutic strategies to improve the often poor prognosis of individuals with peripheral and central nervous system (CNS) injuries. To gain better insight into this complex biological process, a number of large-scale studies have identified candidate genes whose change in expression level could affect axonal regeneration, constituting regeneration-associated genes (RAGs) (Chandran et al., 2016). Indeed, a number of genes when overexpressed or inhibited affect regeneration including transcription factors, inflammatory signals, adhesion molecules and neurotrophins (for a detail review see (Mar et al., 2014)). Such studies provide one strategy toward identifying the molecular control elements of axon regeneration. To uncover additional regulatory genes, a comparison of inbred mouse strains that differ genetically in their regenerative capacity can reveal adaptive evolutionary strategies and the genes involved that favor peripheral or CNS regeneration (Omura et al., 2015).

Dorsal root ganglion (DRG) neurons are located at the interface of the central and peripheral nervous system (PNS). They display an ability to regenerate after peripheral axotomy that is associated with the growth permissive environment of the PNS, which differs from the inhibitory extrinsic cues in the CNS (Hoffman, 2010). Successful regeneration in the PNS also involves induction of an intrinsic growth capacity after axonal injury which is associated with multiple transcriptional changes (Tedeschi, 2011) and can be enhanced by a preconditioning injury, such as a prior nerve crush (Neumann and Woolf, 1999; Lu P et al., 2004; Cafferty, 2004) Similar injury-induced transcriptional changes and enhancement of an intrinsic regenerative capacity is achieved by the dissection related axotomy and dissociation of primary DRGs for the purpose of culturing (Saijilafu and Zhou, 2012; Smith and Skene, 1997), which serves as a “preconditioning” injury relative to a subsequent *in vitro* axotomy (IVA)

DRG neurons comprise a diversity of cell types that can be categorized according to their conduction properties, peripheral and central innervation patterns, morphologies (Basbaum et al., 2009; Abraira and Ginty, 2013) and by the functional modalities they mediate. Recently, classification schemes of different somatosensory neuron subtypes based on the sets of genes they express have revealed a greater heterogeneity than previously appreciated (Chiu et al., 2014; Li et al., 2016; Usoskin et al., 2014). How these multiple different cell types respond to nerve injury and collectively coordinate a regenerative response is unknown.

To understand the molecular changes promoting regeneration, we have taken advantage of two mouse strains C57BL/6 and CAST/Ei, that differ greatly in their regenerative capacity, as measured by regrowth of peripheral sensory neurons on inhibitory substrates both with and without a preconditioning injury (Omura et al., 2015). The CAST/Ei strain shows enhanced axonal outgrowth *in vitro* and a much stronger regenerative phenotype *in vivo,* especially in response to a preconditioning lesion. Here we identify *Ascl1* as a contributor to the improved regenerative phenotype of the CAST/Ei strain and show that the IVA model recapitulates the phenotypic and molecular differences observed in both strains in earlier preconditioning and CNS regeneration studies. We also show that *Ascl1* levels are differentially regulated in the two strains transcriptionally via its promoter activity and post-transcriptionally via the microRNA, miR-7048-3p. Single cell RNAseq of DRG neurons partitioned the cells into three populations: (a) those that expressed genes related to mature neuronal function; (b) those that expressed genes related to cellular differentiation, development and neurogenesis; (c) those that expressed genes related to both of these processes. Cells expressing genes related to cellular differentiation also more abundantly expressed genes associated with a neural-crest like identity as well as stress markers. After injury, the proportion of cells with an immature neural-crest like expression pattern increased more in the C57BL/6 than in the CAST/Ei strain. These immature, neural-crest like cells express Ascl1 target genes at much lower levels than the cells with a mature profile and this may be linked to the poor regenerative outcome.

## Results

### *Ascl1* Mrna is upregulated after injury and its upregulation enhances regeneration

To identify genes responsible for the enhanced regenerative phenotype of CAST/Ei mice, we performed a comparative transcriptional analysis by RNA sequencing DRGs from four different mice strains: C57BL/6, CAST/Ei, DBA/2 and PWK/Ph. These were chosen because of their varied regenerative capacity. The C57BL/6 strain is a poor regenerator, CAST/Ei is the best regenerator and the DBA/2 strain has an intermediate phenotype (Omura et al., 2015). The PWK/Ph strain is evolutionarily distant from the other three strains. Lumbar (L4 and L5) DRGs were harvested from naïve animals and from preconditioned animals with a sciatic nerve crush injury 5 days prior to harvest. The transcriptional profiles obtained clustered according to treatment and strain (Fig. S1).

We focused on the C57BL/6 and CAST/Ei strains, the worst and best regenerators respectively. Comparing the expression level of genes in each strain, we observed that genes upregulated after injury mostly related to an immune response, whereas genes downregulated in the corresponding comparison related to ion transport. To identify the genes associated specifically with the CAST/Ei improved regenerative phenotype, we picked genes whose expression was specifically higher in DRGs of CAST/Ei after injury. Out of the 15,743 genes expressed in our dataset, 696 fit this relative expression pattern. Of these genes, 12 were upregulated at least 16 fold (Table I). The gene with the greatest fold increase after injury in the CAST/Ei strain was *Ascl1* (Table I and Fig. 1A). This gene was also upregulated after injury, in the DBA/2 strain, which also displays an enhanced regenerative phenotype relative to the C57BL/6 strain. Given the high and specific upregulation of *Ascl1* in the CAST/Ei after injury, we focused on the role of this gene in regeneration. ASCL1 is a basic helix-loop-helix transcription factor that plays a key role in cell cycle exit and pro-neuronal differentiation (Castro et al., 2006; Farah et al., 2000; Nakada et al., 2004). The CAST/Ei increased expression in response to injury occurred in neurons (Fig. S2)

**Fig. 1.**
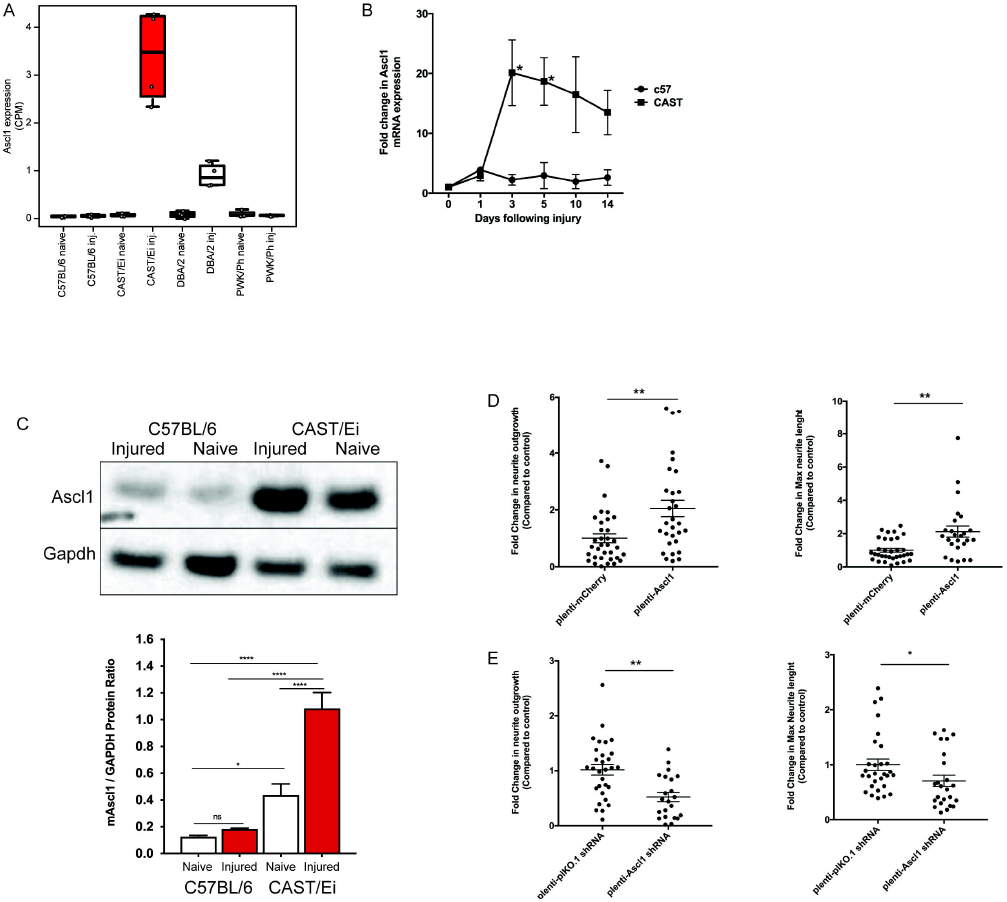
*Ascl1* enhances the neurite outgrowth of CAST/Ei DRG neurons. (A) *Ascl1* mRNA is specifically upregulated in CAST/Ei DRG neurons after *in vivo* injury as assayed by RNAseq (log_2_ (FC) vs CAST/Ei naïve = 5.6, FDR = 2.45E-48). (B) *Ascl1* mRNA levels were significantly upregulated 3 and 5 days post sciatic nerve crush injury and levels were different until day 14 as measured by qRT-PCR. Values are represented as mean ± SEM. [* p< 0.05, Student’s t-test]. (C) The protein expression of Ascl1 is higher in CAST/Ei DRGs than in the C57BL/6 DRGs after *in vivo* sciatic nerve crush injury as shown by western blot and quantified as mean ± SEM of the western blot quantification. [* p< 0.05, * * p< 0.01, One-way ANOVA, Tukey’s post-hoc test]. (D) Forced overexpression of Ascl1 for 7 days in cultured C57BL/6 DRG neurons using pLenti-Ascl1-GFP produced enhanced neurite outgrowth after replating (measured 24 h after replating). Total neurite outgrowth per well as well as maximum neurite length measured by metaexpress showed significant increase in presence of pLenti-Ascl1 compared to pLenti-mcherry (control plasmid). (E) Ascl1 knockdown showed a decrease in total neurite outgrowth and length of longest growing axons compared to control plasmid. Values are represented as mean ± SEM. [* p< 0.05, * * p< 0.01, Student’s t-test].

**Table I.** Genes upregulated at least 16 folds specifically in the CAST/Ei DRG neurons after the preconditioning injury.

| Gene Name | Log2(Fold change) | Corrected p-value |
| --- | --- | --- |
| *Ascl1* | 5.55 | 4.36E-46 |
| *Ifit1* | 5.34 | 8.29E-73 |
| *Ifi44* | 5.19 | 1.18E-58 |
| *Usp18* | 5.19 | 7.94E-72 |
| *Irf7* | 5.08 | 3.02E-90 |
| *Irg1* | 5.06 | 2.53E-07 |
| *Klk4* | 4.86 | 2.29E-88 |
| *Saa3* | 4.63 | 2.33E-10 |
| *Cpz* | 4.52 | 3.94E-23 |
| *Ifit2* | 4.30 | 4.81E-75 |
| *Isg15* | 4.26 | 4.24E-61 |
| *Ifit3* | 4.08 | 1.94E-41 |

We independently confirmed *Ascl1* upregulation in the CAST/Ei strain by qPCR. *Ascl1* mRNA was upregulated within three days after *in vivo* sciatic nerve crush injury in the CAST/Ei DRG and its levels were significantly higher than those observed in C57BL/6 DRGs on day 5 (Fig. 1B). Consistent with the RNAseq and the qPCR results, protein levels of ASCL1 were higher in CAST/Ei injured DRGs compared to either naïve CAST/Ei or injured C57BL/6 DRGs (Fig. 1C). ASCL1 protein levels in naïve DRGs were higher in the CAST/Ei strain than in the C57BL/6 strain, something that we did not observe at the mRNA level (Fig. 1B and Fig. 1C).

To test if *Ascl1’s* higher expression contributed to the improved regeneration, we infected C57BL/6 DRG neurons in culture with an *Ascl1* expressing lentivirus. The infection increased the Ascl1 mRNA five fold (Fig. S2). Following the overexpression we measured neurite outgrowth as a proxy for regeneration. ASCL1 overexpressing neurons showed a significant (p< 0.01) 1.5 fold increase in neurite length (Fig. 1D). Inversely, knockdown of ASCL1 in CAST/Ei neurons infected with a lentivirus expressing an Ascl1 shRNA decreased Ascl1 mRNA levels (Fig. S2), neurite outgrowth (p< 0.01) and maximum neurite length (p< 0.05) compared to neurons infected with a lentivirus expressing a control shRNA (Fig. 1E). These results suggested that at least part of the improved regenerative phenotype observed in the CAST/Ei strain is likely attributable to increased ASCL1 expression.

### The IVA model recapitulates the physiological and molecular phenotype of an *in vivo* preconditioning injury

The culture-and-replate *in vitro* axotomy (IVA) protocol recapitulates many of the biochemical and morphological features of an *in vivo* preconditioning injury (Liu et al., 2013; Saijilafu et al., 2013; Saijilafu and Zhou, 2012; Smith and Skene, 1997). We investigated whether this was also the case for the enhanced regenerative ability of the CAST/Ei DRG neurons over that of the C57BL/6. We grew dissociated DRG neurons for 48h on plastic before replating them in an IVA model on poly-D-lysine and laminin coated coverslips, a substrate permissive to regeneration. We compared the regrowth of these replated neurons to that of neurons plated on poly-D-lysine and laminin immediately after dissociation (“naïve"). Sixteen hours after re-plating the IVA neurons showed extensive long neurites with greater arborization when compared to non-replated naïve neurons (Fig. 2A). In both naïve and IVA neurons, the extent of neurite outgrowth in the CAST/Ei strain exceeded that observed in C57BL/6 sensory neurons. In the naïve state, mean neurite length, measured as the sum of the neurite lengths on each neuron, was 1.6 times greater in CAST/Ei than C57BL/6 (p-value = 4e-4, Fig. 2B). In IVA neurons CAST/Ei neurites were 1.4 times longer than those of C57BL/6 (p-value = 3.7e-3 Fig. 2B). A Sholl analysis revealed that the complexity of the neurites was also greater in the CAST/Ei samples than in the C57BL/6 samples (Fig. 2C). The number of neurites emerging from the cell body (neurite initiation) was similar in both strains, as indicated by the overlapping of the Sholl analysis curves at short distances.

**Fig. 2.**
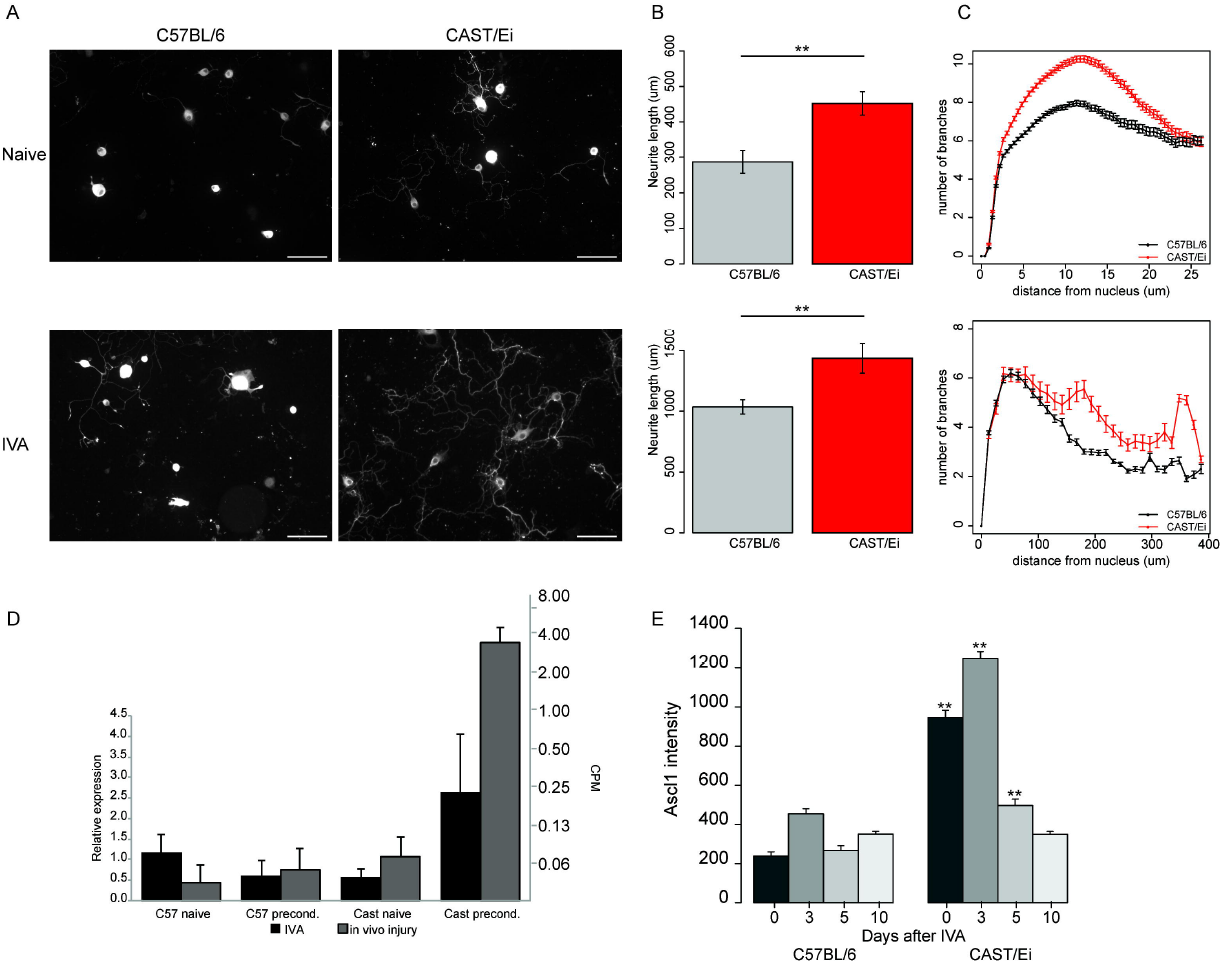
In vitro axotomy recapitulates the phenotypic strain differences in regeneration observed after *in vivo* injury. (A) Representative fluorescence images of C57BL/6 and CAST/Ei DRG neurons stained against TUBB3 highlighting neurite outgrowth in both strains with or without an IVA. [Scale bar = 100 μm] (B-C) Quantification of neurite length (B) and complexity (C) in the naïve (upper panel) and IVA (lower panel) C57BL/6 (black) and CAST/Ei (grey) neurons. The CAST/Ei neurites are longer and more complex with a greater arborization, in both conditions. (D) IVA (black) recapitulates the increase in *Ascl1* mRNA levels observed after in vivo injury (grey). (E) Time course of Ascl1 protein expression levels in C57BL/6 and CAST/Ei neurons using immunocytochemistry. Ascl1 levels were higher up to 5 days after IVA, in the CAST/Ei neurons than C57BL/6 neurons. By day 10, the levels were not different from baseline in both strains after IVA. (* * p≤ 1e-2, Student’s t-test).

We tested if the increase in *Ascl1* mRNA expression observed after an *in vivo* peripheral nerve injury was also observed after IVA. The levels of *Ascl1* mRNA measured by qPCR were upregulated 4.6 fold after IVA in the CAST/Ei strain, consistent with the results obtained from the *in vivo* injury (Fig. 2D). Relative ASCL1 protein levels in neurons measured by quantifying microscopy images (see. Sup. Methods) was higher in CAST/Ei naïve samples than in either the naïve or IVA C57BL/6 samples (Fig. 2E), consistent with the *in vivo* findings. ASCL1 protein levels were further increased three days after IVA in CAST/Ei neurons compared to naïve neurons, consistent with the patterns of mRNA and protein upregulation *in vivo* (Fig. 1). Taken together, these results show that IVA recapitulates both the phenotypic growth and molecular changes in ASCL1 expression observed after an *in vivo* preconditioning injury.

### *Ascl1* is differentially regulated in C57BL/6 and CAST/Ei strains at the transcriptional and at the post-transcriptional levels by miR-7048-3p

The differential regulation of Ascl1 in the two mouse strains could result from a differential transcriptional or post-transcriptional regulation, or both. To gain insight into Ascl1 transcriptional regulation we developed a luciferase reporter assay by cloning either the C57BL/6 or the CAST/Ei promoter of *Ascl1* upstream of the firefly luciferase gene. In both 293T and NIH/3T3 cells, the CAST/Ei promoter of *Ascl1* had stronger basal activity than that of the C57BL/6 promoter (Fig. 3A). We also sought to determine whether post-transcriptional microRNA-mediated regulation of Ascl1 occurred. We sequenced small RNAs of C57BL/6 and CAST/Ei DRGs before and after an *in vivo* preconditioning injury. Of the 1908 annotated murine mature miRNAs deposited in miRBase (release 20), 1123 were expressed in our dataset. Comparing miRNA expression across strains after the same treatment (C57BL/6 naïve vs CAST/Ei naïve and C57BL/6 injured vs CAST/Ei injured), 255 miRNAs were differentially expressed (Fig. 3B, top left panel). When comparing treatments within the same strain (C57BL/6 naïve vs C57BL/6 injured and CAST/Ei naïve vs CAST/Ei injured), 112 miRNAs were differentially expressed (Fig. 3B, top right panel).

**Fig. 3.**
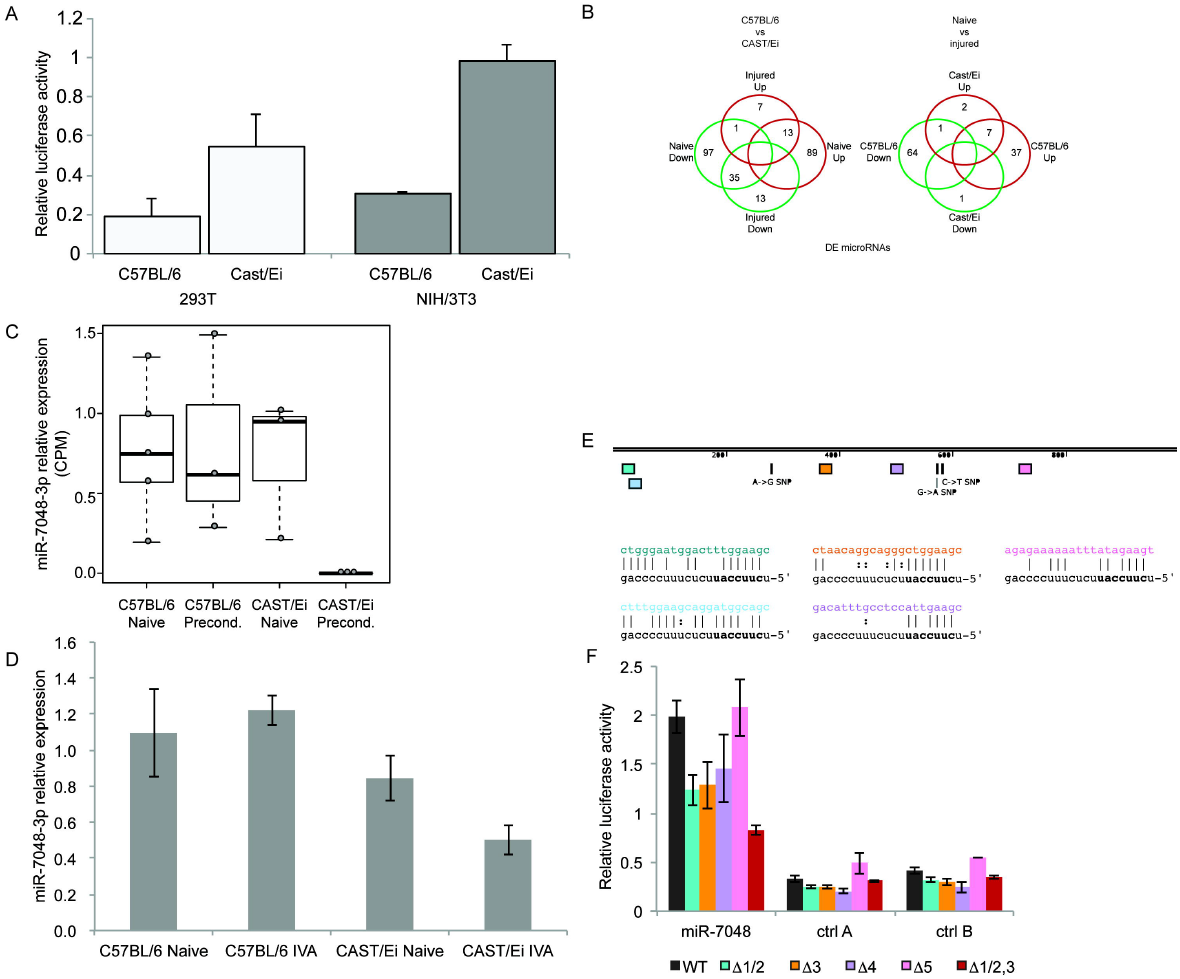
*Ascl1* is differentially regulated in the C57BL/6 and the CAST/Ei strains both transcriptionally and post-transcriptionally. (A) Relative luciferase activity of the C57BL/6 and CAST/Ei promoter of *Ascl1*. The basal activity of the CAST/Ei promoter is higher than that of the C57BL/6 promoter in both 293T and NIH/3T3 cells. The error bars represent the standard deviation (SD). (B) Strain differences are more robust than injury responses in terms of their impact on miRNA expression. 255 miRNAs are differentially expressed when strains are compared (left Venn diagram) whereas only 112 are differentially expressed when treatments (naïve vs injured) are compared (right Venn diagram). (C) miR-7048-3p expression is significantly decreased in the *in vivo* preconditioned CAST/Ei samples compared to either the naïve or preconditioned C57BL/6 samples assayed by small RNAseq. (D) IVA recapitulates the decrease in miR-7048-3p observed in the CAST/Ei after *in vivo* injury. (E) miR-7048-3p has 5 putative binding sites in the 3’UTR of *Ascl1*, two of which (teal and blue) overlap and are positioned in the 5’ extremity of the *Ascl1* 3’UTR. None of the putative binding sites overlap with any of the 5 known genomic differences (represented by vertical bars and annotated with their nucleotide changes) between the two strains. Below is the sequence of the binding sites for each of the putative miR-7048-3p binding sites. (F) Co-transfection of a miR-7048-3p inhibitor and a luciferase reporter harboring the 3’UTR of *Ascl1* in Neuro2A cells increases the luciferase activity over 5-fold when compared to two negative control miRNA inhibitors. The mutation of any of the predicted binding sites but of site #5 decreases the luciferase activity. The error bars represent the standard error of the mean (SEM) of biological replicates.

We screened for miRNAs whose expression decreased after peripheral nerve injury in the CAST/Ei strain only, as a potential explanation for the increased ASCL1 expression specific to this strain in this condition. Out of the 1123 mature miRNAs expressed in the sequenced samples, surprisingly only one displayed a selective decrease after injury in CAST/Ei DRGs: miR-7048-3p (Fig. 3C and D). So far, the existence of this miRNA has only been reported in the mouse genome.

Remarkably, miR-7048-3p has five putative binding sites in the 3’UTR of *Ascl1* in both the C57BL/6 and the CAST/Ei strains. Two of the putative binding sites overlap at the 52’ end of the 3’UTR of *Ascl1* (Fig. 3E). The extended base pairing, the presence of five sites (sites 1 and 2 overlap) and their location in the extremity of the 3’UTR together suggest a functional target (Grimson et al., 2007). To test the effectiveness of miR-7048-3p as a regulator of ASCL1, we cloned the 3’UTR of *Ascl1* downstream of a luciferase reporter and tested its activity. We found that when miR-7048-3p inhibitors were cotransfected with the reporter in the Neuro2A cell line, relative luciferase activity was strongly increased compared to the cotransfection with either of two negative control miRNA inhibitors (Fig. 3F). We then tested if the increase in luciferase activity observed after transfecting miR-7048-3p inhibitor was a direct consequence of the binding of miR-7048-3p to the predicted binding sites on the 3’UTR of the reporter by mutating the predicted binding sites. We abolished the putative pairing of miR-7048-3p to the reporter by replacing the nucleotides predicted to bind positions 2 to 5 of the miRNA with their complementary nucleotide. We abolished binding at each of the five predicted binding sites individually with sites 1 and 2 being treated as a single site due to their overlap. The co-transfection of a miR-7048-3p inhibitor with any of the mutated reporters, except mutated site 5, resulted in a decrease in luciferase activity compared to the wild type reporter (Fig. 3F). Destroying the pairing simultaneously in all three of the most 5’ binding sites further decreased luciferase activity relative to destroying sites 1/2 and site 3 individually (Fig. 3F, Δ 1/2,3). None of the mutations had any effect on the luciferase activity when control miRNA inhibitors were cotransfected. We next investigated whether miR-7048-3p repressed Ascl1 by degrading its mRNA or inhibiting its translation. C17.2 cells transfected with miR-7048-3p inhibitors showed an increase in Ascl1 mRNA expression (Fig. S3) indicating that the repression is via transcript degradation. This effect was specific to Ascl1 as none of the other tested genes showed upregulation after transfection of the miR-7048-3p inhibitor. Collectively, these results show that miR-7048-3p regulates the expression level of ASCL1, and this implies a role for injury-induced decreases in miR-7048-3p in axonal regeneration.

We tested the effect of miR-7048-3p on the regeneration of primary sensory neurons in culture. We inhibited miR-7048-3p in naïve C57BL/6 DRG sensory neurons by nucleofecting a miR-7048-3p inhibitor. This resulted in an increase in total neurite outgrowth (p< 0.01), mean number of processes per neuron (p< 0.01) and an increased complexity of neurites (p< 0.01) compared to transfection with a control miRNA inhibitor (Fig. 4). Consistent with such a neuronal function for miR-7048-3p, a GO term analysis of its predicted targets (see Methods) revealed that of the 84 GO terms in the “biological processes” category that were significantly enriched in the miR-7048-3p putative targets, 18 (21%) were associated with the nervous system (Table SI).

**Fig. 4.**
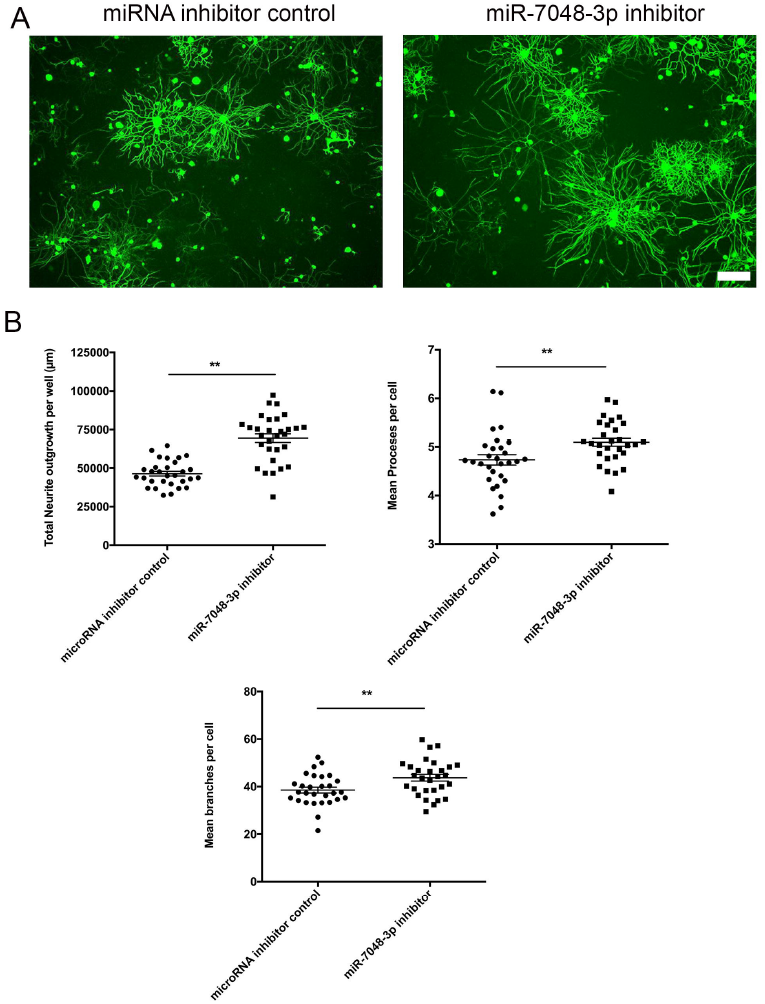
miR-7048-3p enhances axonal outgrowth. (A) Representative images of Tubb3 stained DRG sensory neurons following transfection with miRNA inhibitor control or miR-7048-3p inhibitor. (B) miR-7048-3p inhibitor showed significant increase in total neurite outgrowth, number of processes per cell and mean branches per cell. Values are represented as mean ± SEM. [* * p< 0.01, Student’s t-test, Scale bar = 100 μm].

### Single cell RNAseq defines three DRG neurons populations and the CAST/Ei response to injury

Transcriptional analysis of mixed cell populations obscures processes that operate at the single cell level, especially in the context of cellular responses to the environment, such as injury, which are known to differ among neuronal subtypes (Hu et al., 2016). We performed single cell RNA sequencing (scRNAseq) of naïve and IVA C57BL/6 and CAST/Ei DRG neurons using a microfluidics chip (Fluidigm C_1_) that selectively captures cells greater than 17um in diameter, thus minimizing glial contamination. We confirmed neuronal capture by visual inspection of the capture sites and excluding from the analysis cells with morphologies inconsistent with DRG neurons (Fig. S4). We confirmed that, for each of the sequenced cells, Tubb3 expression was detected (Fig. 5A).

**Fig. 5.**
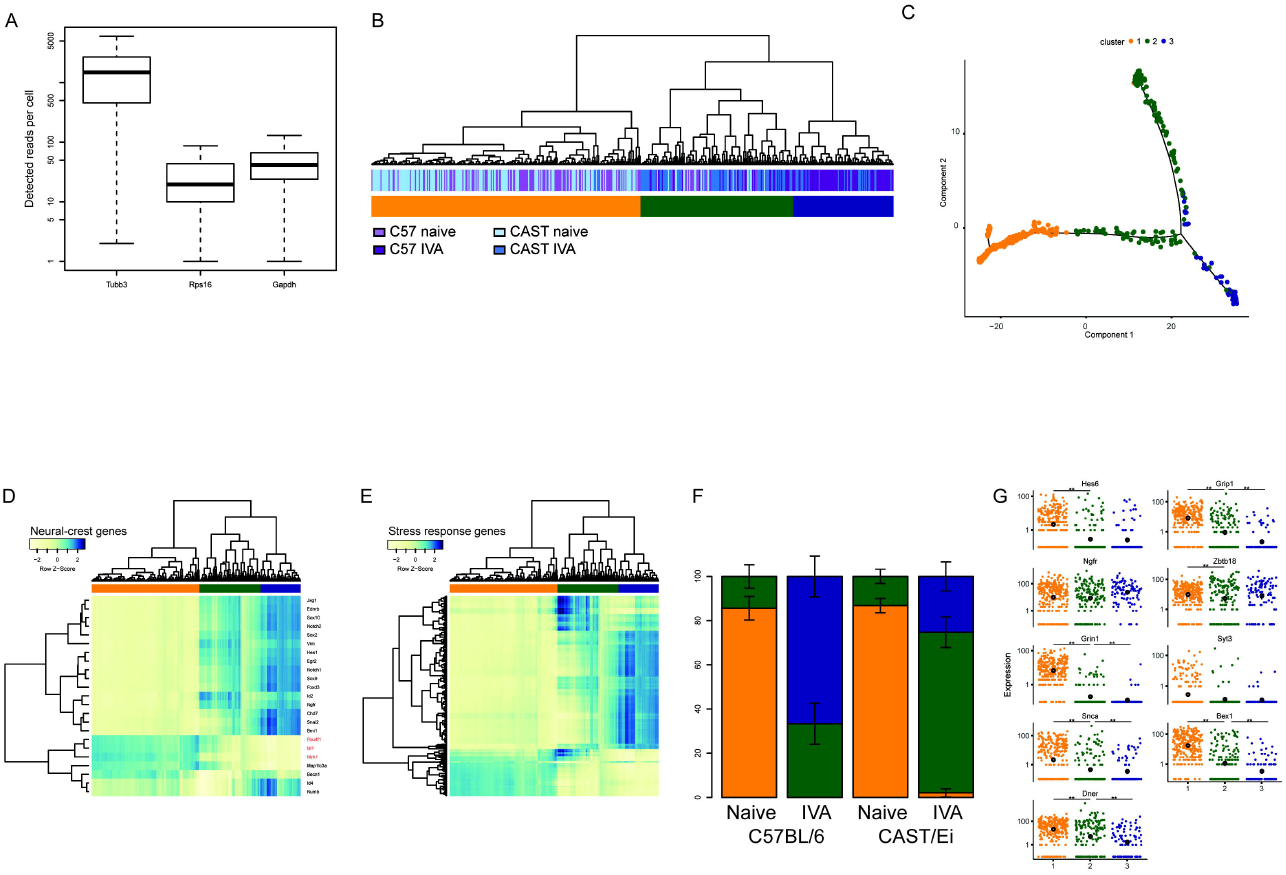
scSeq highlights the existence of two distinct clusters of DRG neurons. (A) Absolute read count detected for Tubb3, Rps16 and Gapdh showing that Tubb3 is detected in each of the sequenced cell, confirming their neuronal identity. (B) Hierarchical clustering of the single cell RNAseq sample according to the expression of the 2,000 genes contributing the most to the transcriptional profile. Three broad clusters (color coded in orange, green, and blue) are identifiable. (C) Result of chronologically ordering the cells along pseudotime using the top 2000 genes identified. The orange (mostly naive) cells mark the beginning of the pseudotime course. The cells belonging to the green cluster are found at the interface between those belonging to the orange and the blue clusters. (D) Heatmap representation of the expression of genes related to neural crest. The three genes (Isl1, Pouf1 and Ntrk1) that are turned on when cells commit to differentiation and exit cell cycle are highlighted in red. (E) Heatmap representation of the expression of the genes annotated as “Cellular response to stress”. (F) Proportion of cells found in each of the two groups identified in A for each of the sample type; color-coded as in A. (G) Expression level of nine expressed Ascl1 target genes as assessed by scRNAseq showing that Ascl1 targets are preferentially expressed in the cells of the orange and green clusters. The large dot in each group represents the trend expression level of this group.

We selected the top 2000 genes with the greatest contribution to the transcriptional profile (see Supp. Methods) and used this gene set to perform unsupervised hierarchical clustering. The cells clustered into two large groups based on treatment, naive versus IVA, as observed in the bulk RNA sequencing (Fig S1). At the next level, the cells clustered into three groups, designated orange, green and blue. The orange group contained almost exclusively naive cells, the blue group contained exclusively IVA cells and the green group contained a mixture of naive and IVA cells (Fig. 5B). To assess the extent to which this unbiased clustering might parallel the distribution of known DRG neurons markers, we performed a new hierarchical clustering with a set of known markers (Chiu et al., 2014) for four DRG neuron types: proprioception (color coded in black), thermoception (pink), pruriception (orange) and tactile function (purple). The clustering of the neurons into three groups was globally preserved when clustered by this gene subset (Fig. S5). This clustering did not offer any insight into the biology of DRG neurons because specific cell types did not associate with regeneration.

We performed a differential expression analysis of the cells in each cluster to identify their defining genes, i.e. the genes more abundantly expressed in one group compared to the other. The defining genes of cells in the orange vs. the blue or green clusters related to mature neuronal processes (ion transport, neurological processes, synaptic transmission, behavior, etc.) (Fig. S6). The genes enriched in the cells of the blue cluster compared to either the orange or the green cluster highlighted processes related to cell division (negative regulation of cellular processes, cell division, nuclear division, etc.). The genes more abundant in the green cluster highlighted an intermediate expression profile. This intermediate profile suggested a chronology of cell identities in response to IVA where neurons transition from the orange to the green and finally to the blue cluster. We tested this suggestion by performing a monocle2 analysis (Trapnell et al., 2014) on the single cell sequencing data ordering the cells along a pseudotime trajectory. As hypothesized, the cells of the green cluster were observed at the transition between the cells of the orange and the blue clusters (Fig. 5C). This analysis also confirmed the existence of two groups of naive cells. A major one (the orange one), at the beginning of the trajectory and a minor one (the green one) observed at a later pseudotime.

Because the DRG develops from the neural crest, we tested the expression of neural crest markers in our single cell data. Of the 23 genes documented to be associated with neural crest cells (Gilbert, 2000; Lee et al., 2007; Li et al., 2007), and expressed in our dataset, 15 were upregulated in the cells of the blue cluster compared to the cells of the orange cluster. Among this set of upregulated genes in the blue cluster were markers of boundary cap cells, Egr2 (aka Krox20) (Hjerling-Leffler et al., 2005) and Sox10. Among the differentially expressed genes more abundantly expressed in the cells of the orange cluster were cell type-specific markers known as terminal selectors (Hobert, 2008) such as Isl1 (aka Islet1), Pou4f1 (aka Brn3a) and Ntrk1 (aka TrkA) (Lallemend and Ernfors, 2012) (Fig. 5D). These genes were expressed at intermediate levels in the cells of the green cluster (q-value < 1e-4), further supporting the positioning of these cells as intermediate on the injury response of DRG neurons. Genes annotated with the “cellular response to stress” GO terms were more abundantly expressed in the cells of the blue cluster (Fig. 5E). These genes included Hsp90b1, a heat shock protein, Rad18, an E3 Ubiquitin ligase implicated in the response to UV damage, and several genes related to ER stress such as Atf4 and Nfe2l2.

In the naive state, the proportion of C57BL/6 and CAST/Ei cells in each of the three clusters did not significantly differ (Fig. 5F, p-value C57BL/6 naive vs CAST/Ei naive = 0.5). After IVA, the proportion of cells found in the green and blue clusters increased significantly in both strains but with different magnitude. In the C57BL/6, 33% of the IVA cells are found in the green cluster and 67% of the cells in the blue cluster (Fig. 5F). In contrast, CAST/Ei neurons were predominantly found in the green cluster (73%), the intermediate step between the orange and blue cluster. Further, a small but significant proportion of CAST/Ei neurons were found in the orange cluster after IVA (2%, p-value between C57BL/6 and CAST/Ei = 0.018). Thus cells that expressed genes related to neurogenesis and stress preferentially emerged in the C57BL/6 genotype after IVA. These observations suggested differences in the proportion of cells in each cluster could account for the observed differential regeneration phenotypes.

Some neural crest markers were expressed in the naïve condition in both the single cell and whole DRG RNAseq. 37 genes associated with neural crest cells were expressed in the bulk RNAseq dataset. In C57BL/6, 12; in CAST/Ei 10; in DBA/2 15; and in PWK/Ph 10 of these genes were significantly upregulated after *in vivo* peripheral nerve injury. Immunocytochemistry against Nestin and Msi2 showed an increase of these two neuronal precursor markers after IVA in both C57BL/6 and CAST/Ei (Fig. S7) showing that, neurogenesis-related gene upregulation is a general response to injury.

The change in abundance of each cell population following nerve injury raised the possibility that both the *in vivo* injury and the IVA induced transdifferentiation of the neurons a phenomenon that occurs in Schwann cells (Baggiolini et al., 2015; Motohashi and Kunisada, 2015). We observed BrdU+ cells among C57BL/6 DRG neurons (TUBB3+ cells) suggesting DNA synthesis in S phase of the cell cycle, but none of the neurons observed had the characteristic nuclear envelope breakdown associated with M phase (Fig. 6). This observation suggested that some cells entered the cell cycle aberrantly without completing cell division. This finding is consistent with data showing that after an insult associated with a chemotherapeutic agent, DRG neurons can re-enter the cell cycle but do not complete it and get shunted toward apoptosis (Fischer et al., 2001). Thus injured neurons may have atavistic properties that results in regression to a dedifferentiated state in the absence of cell division and that neurons in this state are less competent regenerators.

**Fig. 6.**
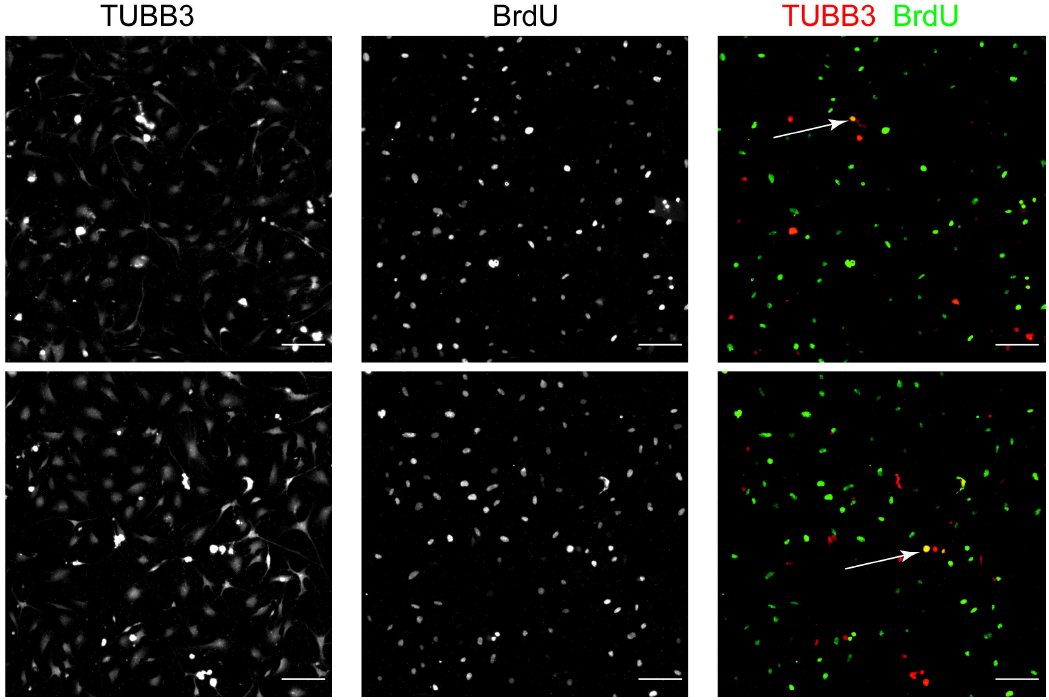
BrdU incorporation in sensory neurons. Some, but not all, of the dissociated DRG neurons grown for 4 days in culture in presence of BrdU are stained for both TUBB3 and BrdU (highlighted by the white arrows). [Scale bar = 100 μm]

### Ascl1 target genes are associated with enhanced regeneration in CAST/Ei mice

Our observations suggest that the enhanced regeneration phenotype of CAST/Ei neurons is associated with retention after injury of a terminally differentiated neuronal identity, i.e. cells in the orange cluster, while a reduced regenerative capacity may reflect conversion to a de-differentiated profile that also exhibits stress markers. The higher proportion of orange and green cells after IVA in the CAST/Ei strain than in the C57BL/6 strain suggests that cells of the orange and green clusters are linked to the better regenerative outcome. If so cells of these two clusters, would have a higher expression of genes promoting regeneration, such as Ascl1. Ascl1 was not detected in the scRNAseq data set nor was it detected in other experiments using similar samples and technologies (Hu et al., 2016; Li et al., 2016; Usoskin et al., 2015) thus ruling out a sample collection problem. Although Ascl1 was not detectable in the scRNAseq datasets, the expression of its validated targets is a reasonable proxy for Ascl1’s activity. A list of validated Ascl1 targets (Treutlein et al., 2016; Wapinski et al., 2013) revealed that 7 of the 9 detectable Ascl1 targets were significantly more abundantly expressed in cells of the orange cluster compared to either the cells of the green or blue cluster (Fig. 5G). Of those, 5 were more abundantly expressed in the cells of the green cluster than in the cells of the blue cluster. This suggested that ASCL1’s expression was highest in the cells of the orange and green clusters and lowest in the cells of the blue cluster.

## Discussion

CAST/Ei neurons have a higher axonal regeneration potential compared to several other inbred mouse strains and activin signaling was identified as one key mechanism for this enhanced regenerative capacity (Omura et al., 2015). However, activin administration alone was insufficient to induce a level of regeneration equivalent to that in CAST/Ei mice, indicating involvement of other molecules or pathways. In the present study using bulk and single cell RNAseq before and after axonal injury, we explored which gene expression profiles may contribute to this phenotype. We identified Ascl1 as one of the most differentially and selectively regulated transcription factors in CAST/Ei DRGs after peripheral nerve injury. Forced expression of Ascl1 in C57BL/6 DRG neurons enhanced axonal outgrowth and knockdown of Ascl1 in CAST/Ei DRG neurons decreased axonal growth. Ascl1 is also known to enhance regeneration of retinal ganglionic cells in zebrafish (Ueki et al., 2015), but how Ascl1 enhances regeneration is not known. Using single cell RNAseq we found that Ascl1 target genes associated with enhanced regeneration potential are expressed at higher levels in CAST/Ei neurons after injury. Consistent with these findings Ascl1 knockdown decreases Gap-43 expression and axonal regrowth after spinal cord injury (Williams et al., 2015).

Ascl1 is an essential transcription factor for the transdifferentiation of fibroblasts into neurons (Vierbuchen et al., 2010; Wainger et al., 2015) and regulates genes responsible for neuronal differentiation and axon guidance, such as epherins, semaphorins, Dcc and Slit (Borromeo et al., 2014). Regulation of axon guidance molecules that contribute to growth cone extension may contribute to the Ascl1-associated regeneration program. Ascl1 also regulates pathways essential for neuronal differentiation, such as Wnt signaling which has been implicated in cardiac regeneration (Ozhan and Weidinger, 2015). Further studies are required to identify the target genes regulated by Ascl1 in sensory neurons and associated with regeneration.

To explore profiles of individual sensory neuron populations associated with the differential strain response to axonal injury we performed single cell RNAseq on DRG neurons. High level clustering first distinguished cells according to treatment but then clearly distinguished three populations that we designated orange, green and blue. The genes driving this partition were related to neurogenesis and cell cycle in the blue cluster, and a mature neuronal phenotype in the orange cluster. The presence of many cells expressing neurogenesis markers, including those of the neural crest, in mature postmitotic DRG neurons may reflect a transdifferentiation-like phenomenon resulting from axonal injury. Most notable was that within this context, the preconditioning injury evoked a highly differential population level response in the two strains. Following IVA, more C57BL/6 DRG neurons shifted to the blue cluster than did the CAST/Ei DRG neurons. The features of cells in the blue cluster resembled to some extent transdifferentiation. In addition to markers of neurogenesis these cells also exhibited features that suggested an aberrant attempt to re-enter the cell cycle as indicated by some C57BL/6 cells with positive BrdU labeling. A greater tendency to aberrant cell cycle entry and a stress response may be less conducive to implementing an effective regeneration program. A stress response in the context of axotomy induced injury is known to be associated with a poor regenerative output (Park et al., 2008). The increased expression of Ascl1 in CAST/Ei after *in vivo* injury may contribute to maintaining a more differentiated state and thereby permit more effective regeneration. Further, the pseudotemporal ordering of the cells revealed the presence of an intermediate cellular type, the cells of the green cluster. Neurons from the CAST/Ei strain preferentially adopt this cell type rather than the type of the blue cluster observed in the C57BL/6 neurons. Thus, the cellular identity of the CAST/Ei neurons is less affected by IVA than it is in the C57BL/6 neurons. This decrease magnitude in response is likely linked to the improved regenerative phenotype of the CAST/Ei neurons. The finding that miR-7038-3p targets Ascl1 and shows a selective decrease after injury in CAST/Ei DRGs suggests that other miR-7038-3p targets may be involved in the enhanced regenerative capacity of CAST/Ei DRGs.

Interestingly, although the single cell RNAseq revealed the emergence of a population of cells after IVA, it did not clearly separate the cells based on strain as the bulk RNAseq did. Within the overriding blue-orange-green clusters we could not readily distinguish further sub-clusters. These results demonstrate how single cell analyses can reveal population behaviors that are not apparent when cells are averaged as they are in bulk RNAseq. Several methodological differences may explain these clustering patterns. First, non-neuronal cells make up the vast majority of DRG and are included in the bulk transcriptomes whereas the single cell analysis included only neurons. Thus non-neuronal cells, particularly glial cells may contribute to the observed clustering in the bulk RNAseq, whereas the profile derived from single cell RNAseq pertained exclusively to neurons. However, the bulk RNAseq clustering can be recapitulated from the scRNAseq data by *a priori* summing each transcript in subsets of cells from the same condition and thereby creating a synthetic-bulk transcriptome (Wu et al., 2014) (data not shown).

A previous report identified the Inhba mRNA, which encodes the β_A_ subunit of both activin and inhibin proteins, as an enhancer of regeneration responsible for the CAST/Ei regenerative phenotype (Omura et al., 2015). Increasing the levels of any of the three forms of activin (β_A_ β_A_, β_A_ β_B_, or β_A_ β_B_), improved regeneration of the C57BL/6 neurons in both the peripheral and central nervous system. Interestingly, Inhbb, which encodes a different variant of the β subunit (β_B_), was shown to restrain the population of stem cells in the mouse olfactory epitelium via a complex feedback network, which involves Ascl1 (Gokoffski et al., 2011). Our data suggest that regenerative output is positively affected by the maintenance of a differentiated state. Activin’s role in inhibiting stemness could be central to its regenerative role, as we have found for Ascl1.

## Experimental procedures

### Mice

C57BL/6 mice were purchased from Charles River, and CAST/Ei mice from Jackson Laboratory (Bar Harbor). Adult (8-10 weeks old) animals of both sexes were used. DRG neurons were harvested and dissociated as described in (Saijilafu and Zhou, 2012). The left sciatic nerve of adult C57BL/6 and CAST/Ei male mice (8-10 weeks) was exposed at the sciatic notch and crushed with smooth forceps for 30 s. All *in vivo* injuries were performed in accordance with institutional animal care at Boston Children’s Hospital, Harvard Medical School and University of California, Santa Barbara.

### In vitro axotomy

DRG neurons were dissociated as previously described (Saijilafu and Zhou, 2012) and plated on bare plastic at high density (5,000 to 10,000 cells per cm^2^) and allowed to grow for 48 h with the addition of ara-c 24 h after plating. After 48 h, the DRG neurons were lifted using trypsin and lightly triturated to ensure dissociation of all processes.

### Ascl1 infection and neurite outgrowth assay

mCherry and Ascl1 were cloned into pLenti CMV GFP Dissociated sensory neurons were plated on PDL-laminin coated 12 well plates the next day, neurons were transduced with virus. Media was replaced every 48 h and cultures were replated and fixed 24 h after replating. Images were captured and neurite outgrowth was automatically quantified using the MetaXpress software.

### Knockdown of Ascl1

Mission control plasmid containing shRNA sequences for Ascl1 or shRNA control vector, containing a non-specific shRNA, were purchased from Sigma-Aldrich.

### Luciferase reporter assays

*Ascl1 promoter.* The 5kb genomic region upstream of the *Ascl1* start site was amplified from genomic DNA of either C57BL/6 or CAST/Ei. NIH/3T3 cells were plated 24 hours before 72 hours after transfection luciferase activity was assessed

*Ascl1 3’UTR.* The 3’UTR of *Ascl1* was cloned downstream of hLuc in the pEZX-MT06 vector (Genecopoeia). Neuro2A cells were plated and transfected with 100ng of pEZX-MT06-Ascl1 vector and a final concentration of 25nM of the miRNA inhibitor (Exiqon) using Lipofectamine 2000. 48 hours after transfection luciferase activity was assessed

### Single cells RNA sequencing

After harvesting as described above, cells were counted and resuspended at a concentration of 310 cells per ul, stained for viability on ice for 30 minutes and then captured by the C_1_ from Fluidigm using the large chip and processed as described by the manufacturer.

Reads were mapped to the mm10 mouse genome using Bowtie2 (Langmead and Salzberg, 2012), duplicate reads removed using samtools (Li et al., 2009) and the reads summarized on a gene annotation using RsubReads (Gentleman et al., 2004; Liao et al., 2013). The resulting counts per million (cpm) matrix was denoised using singular value decomposition (SVD).

## miRNA target prediction

To identify putative miRNA targets, we obtained from the Ensembl database, all the mouse transcripts for which an orthologue exists in the human genome. We then scanned each 3’UTR sequence against the seed of each miRNA. We defined a site to be a putative target site of a given miRNA if there is at least a 6 nucleotides complementarity between the 3’UTR and the miRNA seed.

### GO term analysis

Transcripts identified in the bulk samples mRNASeq were ranked according to the number of putative miR-7048-3p targets their 3’UTR contains. Enrichment analysis was performed using GOrilla (Eden et al., 2009).

GO term enrichment of the genes identified in the single cell analysis was also performed using GOrilla and visualized using REViGO (Supek et al., 2011)

### miR-7048-3p inhibition in DRG neurons

Dissociated DRG neurons were layered over 6 ml of a 10% BSA solution for gradient separation removal of Schwann cells, myelin and other debris. Purified DRG neurons were transfected with 100 nM miRCURY LNA microRNA inhibitor (mmu-miR-7048-3p, 5nmol, 5`fluorescein labeled; cat # 4106101-011, Exiqon) or inhibitor control (miRCURY LNA™ microRNA inhibitor control, 5nmol, 5`-fluorescein labeled; cat #199006-011) by electroporation in a Nucleofector II interfaced with a Lonza nucleocuvette strip. After 24 hours, cells were fixed and stained. Images were captured at 10X and automatically quantified using the MetaXpress software.

### Statistical methods

Statistical significance of differences among means was evaluated using either Student’s t-test or one-way ANOVA with a Turkey post-hoc test. Differential expression analysis of the bulk sequencing was performed using edgeR (Robinson et al., 2010). whereas monocle 2 (Trapnell et al., 2014) was used for single cell sequencing differential expression analysis.

## Author Contributions

V.L. and K.S.K. planned the project and designed the experiments. V.L., B.S., C.J.W. and K.S.K analyzed the data and wrote the paper. V.L., B.S., M.G., E.G., M.W.P., Y.C., E.H., G.C., and M.C. performed the experiments.

## Accession Numbers

The small RNA sequencing and the single cell sequencing generated in this study have been released to GEO under accession GSE98417.

## Acknowledgments

The authors thank Sara Solla and Yasser Roudi for useful discussion and acknowledge the use of the NRI-MCDB Microscopy Facility, funded by NIH Grant No. 1S10OD010610-01A1. The authors would like to thanks Jenny Shao for the technical support and Dr. Lee Barrett for his help with imaging at Assay Development Screening Core (ADSF), funded by IDDRC grant (NIH/NICHD, P30 HD018655) at Boston Children’s Hospital. V.L. and B.S. are funded by a fellowship from the Canadian Institutes of Health Research. This work was supported by the Dr. Miriam and Sheldon G. Adelson Medical Research Foundation (K.S.K.)

